# Myosin-gelsolin cooperativity in actin filament severing and actomyosin motor activity

**DOI:** 10.1101/2020.09.02.279729

**Authors:** Venukumar Vemula, Tamas Huber, Marko Usaj, Beáta Bugyi, Alf Mansson

## Abstract

Actin is a major intracellular protein with key functions in cellular motility, signalling and structural rearrangements. Its dynamic behavior with actin filaments (F-actin) polymerising and depolymerising in response to intracellular changes, is controlled by actin-binding proteins (ABPs). Gelsolin is one of the most potent filament severing ABPs. However, myosin motors that interact with actin in the presence of ATP also produce actin filament fragmentation through motor induced shearing forces. To test the idea that gelsolin and myosin cooperate in these processes we used the in vitro motility assay, where actin filaments are propelled by surface-adsorbed heavy meromyosin (HMM) motor fragments. This allows studies of both motility and filament dynamics using isolated proteins. Gelsolin (5 nM) at very low [Ca2+] (free [Ca2+] ∼6.8 nM) appreciably enhanced actin filament severing caused by HMM-induced forces at 1 mM [MgATP], an effect that was increased at increased HMM motor density. This finding is consistent with cooperativity between actin filament severing by myosin-induced forces and by gelsolin. As further support of myosin-gelsolin cooperativity we observed reduced sliding velocity of the HMM propelled filaments in the presence of gelsolin. Overall, the results corroborate ideas for cooperative effects between gelsolin-induced alterations in the actin filaments and changes due to myosin motor activity, leading among other effects to enhanced F-actin severing of possible physiological relevance.

## Introduction

Actin is a major cytoskeletal protein, constituting 5 to 10% of the cellular protein content in eukaryotes [1, 2]. It has vital roles in muscle contraction and non-muscular cell motility but also in a variety of other cell functions such as intracellular transport, cytokinesis, membrane dynamics, cell signalling and regulation of cell–cell contacts [3]. In muscle contraction and several other processes, actin filaments (F-actin) interact with the molecular motor myosin II to produce force and motion by sliding of actin and myosin relative to each other. Other functional roles of actin rely on its dynamic properties with polymerisation and depolymerisation regulated by a range of actin-binding proteins (ABPs) and varying cellular conditions [4-7].

Gelsolin is one of the most abundant and potent actin filament severing, capping and nucleating proteins [8-13] among the plethora of ABPs [14, 15] that govern remodelling of the actin cytoskeleton in response to various cues [8]. The gelsolin activity is regulated by the complex interplay between calcium (Ca^2+^), polyphosphoinositide 4, 5-bisphosphate (PIP_2_) and ATP [16], but it is also partially activated by low pH [17] and high temperature (30–40 °C) [18, 19]. Binding of Ca^2+^ to several conserved sites on gelsolin, with Ca^2+^ affinities ranging from the 100 nM to 10 µM, changes gelsolin structure to facilitate binding to actin and subsequent actin filament severing [11, 20] by weakening the noncovalent bonds between actin subunits. After severing, gelsolin remains attached to the barbed end (fast polymerizing plus end) of the filament, thus blocking addition of actin monomers (G-actin). However, gelsolin can also nucleate polymerization by binding to two G-actin units in vitro [19].

The in vitro motility assay is a frequently used tool in studying the motion generated by the interaction between myosin and actin, powered by turnover of MgATP [21, 22]. Generally, myosin motor fragments such as heavy meromyosin (HMM) are adsorbed to modified glass surfaces in a controlled environment while the propulsion of fluorescently labelled actin filaments is observed using microscopy. The speed of the actin filament movement (sliding velocity) depends on several factors [23-25], such as the density of myosin heads (HMM), MgATP concentration, ionic strength, pH, temperature and the surface modification used for HMM adsorption. A lowered HMM surface density leads to reduced actin filament sliding velocity, but also to a reduced filament fragmentation during the HMM propelled sliding [26]. Such filament fragmentation is the basis for an increased number of filaments and reduced average length with time in the in vitro motility assay. However, importantly, actin filament fragmentation due to myosin motor activity may also have critical roles in cellular physiology [7, 27-29] either alone or in cooperation with other ABPs, e.g. as proposed for cofilin [28, 30]. In accordance with evidence that actin filaments exhibit intrinsic structural polymorphism in response to varying environmental conditions [31-33], structural transitions along F-actin induced by ABPs are believed to be important for cooperation with fragmentation due to myosin motor activity [29, 34]. Thus, several studies suggest that binding of a range of ABPs allosterically produce long-range structural transitions along the actin filament [35-44]. For instance, gelsolin binding to an actin filament has been found to produce changes of this type [29, 45, 46], which may be expected to change the mode of actin binding of other proteins including myosin. Furthermore, one may consider the possibility of cooperativity between the filament severing induced by gelsolin binding on the one hand and myosin activity in the presence of MgATP on the other hand [45, 46]. However, to the best of our knowledge, these issues have not been previously studied.

Here, we therefore investigated the effect of gelsolin on HMM-induced actin filament sliding in the in vitro motility assay to assess possible gelsolin-induced changes in the myosin binding to actin or in the subsequent generation of motion. We also studied gelsolin-mediated F-actin severing during HMM propelled actin filament sliding and when actin filaments were bound to surface-immobilized HMM in the absence of motility. The experiments were performed either at a low [Ca^2+^] where little activation of actin binding and severing by gelsolin is expected [47, 48] or at sufficiently high [Ca^2+^] expected to fully activate gelsolin [48]. The results provide evidence for cooperative effects between HMM and gelsolin on the actin filament properties and on actomyosin motor function. Thus, increased gelsolin-induced actin filament severing was observed upon increased HMM surface densities in the in vitro motility assay. Moreover, gelsolin binding to actin, also at very low concentrations of free Ca^2+^, resulted in reduced HMM-induced actin filament sliding velocity, suggesting that gelsolin-binding to an actin filament makes the motion-generating actin-myosin interaction less effective. These effects, in turn, accord with the idea of gelsolin induced long-range structural changes along the actin filament. In the absence of motility, there was a one-to-one ratio between the number of filament severing points along myosin-bound actin filaments and binding of Alexa-647 labelled gelsolin to the filament barbed ends. The situation was different during motility suggesting that the long-range structural changes induced by gelsolin binding lead to cooperative filament severing with myosin only in the presence of ATP and motor activity.

## Materials and Methods

### Ethical statement

Animal handling and experiments were approved by the Regional Ethical Committee for Animal Experiments in Linköping, Sweden (reference number 73-14).

### Chemicals

Rhodamine Phalloidin, NHS-Rhodamine (N-hydroxy-succinimidyl), and Alexa Fluor™647 C_2_ Maleimide (Alexa-647), N-(1-Pyrenyl)iodoacetamide (pyrene) were purchased from Thermo Fisher Scientific, Stockholm, Sweden. Other analytical grade chemicals and reagents used in this study were purchased from Sigma-Aldrich Sweden AB, Stockholm, Sweden unless otherwise stated.

### Protein preparations

Actin was purified from leg and back muscles of rabbit using the acetone powder method and was quickly snap frozen in liquid nitrogen [49], followed by storage at −80 °C. Myosin II was isolated from rabbit leg muscles, and subsequently digested with TLCK-treated α-chymotrypsin to obtain heavy meromyosin (HMM) [50], which was frozen in the presence of 2 mg/ml sucrose and stored at −80 °C. F-actin (0.25 mg/ml) was labelled with rhodamine phalloidin at a molar ratio of 1:1.5 (actin:rhodamine phalloidin) in 10 mM 4-morpholinepropane-sulfonic acid (MOPS) buffer at pH 7.0 containing 60 mM KCl, 2 mM MgCl_2_, 0.1 mM Ethylene glycol-bis(2-aminoethylether)-N,N,N′,N′-tetraacetic acid (EGTA) and 3 mM sodium azide (NaN_3_). NHS-Rhodamine labelling of F-actin was performed according to the manufacturer’s instructions. Protein concentration (HMM, actin) was measured by UV absorbance spectroscopy, whereas protein purity was confirmed by sodium dodecyl sulphate polyacrylamide gel electrophoresis (SDS-PAGE).

The plasmid DNA of His-tagged recombinant full-length human cytoplasmic gelsolin (pET21d (+)) was transformed into *E. coli* BL21 (DE3) cells [51, 52]. A fresh colony of *E. coli* was grown in LB broth at 37 °C until the OD_600_ reached 0.6–0.8 and then induced with 1 mM isopropyl β-D-1-thiogalactopyranoside overnight at 25 °C. The cells were collected by centrifugation (6000 g, 5 min, 4 °C, Sigma 4-16KS tabletop centrifuge), lysed in lysis buffer (5 mM Tris, 300 mM NaCl, 5 mM imidazole, 1 mM ATP, 1 mM phenylmethylsulfonylfluoride, 7 mM β-mercaptoethanol, 30 mg/mL DNase with protease inhibitor cocktail (P8465, Sigma-Aldrich) (pH 8.0)) and then sonicated and ultracentrifuged (440,000 g, 35 min, 4°C; MLA80 rotor, Beckman Optima™ MAX-TL). The supernatant was applied to a Ni-NTA column (Machery-Nagel, Düren, Germany), washed with lysis buffer and eluted with 250 mM imidazole in lysis buffer. Fractions containing gelsolin were dialyzed (20 mM Tris, 1 mM EGTA (pH 8.0)) and further purified on a Source 15Q anion exchange column (GE Healthcare) with the application of 50 mL buffer I (20 mM Tris, 20 mM NaCl, 1 mM EGTA (pH 8.0)), 50 mL buffer II (10 mM Tris, 0.1 mM EGTA (pH 8.0)), 50 mL buffer III (20 mM Tris, 2 mM CaCl_2_ (pH 8.0)) and a 100 mL linear gradient of buffer III and buffer IV (20 mM Tris, 1 M NaCl, 0.1 mM EGTA (pH 8.0)). The gelsolin-containing fractions were dialyzed (5 mM HEPES, 50 mM NaCl, 0.1 mM EGTA (pH 8.0)), and gel filtered on a Superdex 200 column (GE Healthcare) equilibrated with dialysis buffer. Purified gelsolin was collected, concentrated (Vivaspin10K cut-off tubes (Sartorius, Göttingen, Germany); 3000 g, 4 °C, 4-16KS tabletop centrifuge), snap frozen in liquid nitrogen and stored at -80 °C. The protein concentration was measured by spectrophotometry (ε_280_ = 1.29 mL mg^-1^ cm^-1^). Gelsolin was labelled by Alexa-647 (8-fold molar excess) for 2 h at room temperature. The unbound dye was removed by using a PD-10 column (GE Healthcare). The final protein and probe concentrations were determined spectrophotometrically. The molar ratio of the bound probe to gelsolin was 0.67.

### Dilution-induced depolymerisation assays

The G-actin bound Ca^2+^ was replaced to Mg^2+^ by adding 50 μM MgCl_2_ and 200 μM EGTA (final concentrations). Mg^2+^-G-actin in buffer A (4 mM Tris, 0.1 mM CaCl_2_, 0.2 mM ATP, 0.005% NaN_3_, 0.5 mM β-mercaptoethanol (pH 7.8)) at 1 μM concentration (50% pyrene labelled) was polymerized overnight by adding 2 mM MgCl_2_ and 100 mM KCl (final concentrations). The F-actin sample was then diluted to 20 nM with Ca^2+^-free polymerization buffer (4 mM Tris, 0.2 mM ATP, 0.005% NaN_3_, 0.5 mM β-mercaptoethanol, 2 mM MgCl_2_, 100 mM KCl (pH 7.8)) supplemented with 100 μM CaCl_2_ or 1 mM EGTA to obtain the desired Ca^2+^ concentrations. Depolymerisation rates were estimated by linear fitting of the normalized pyrene transient curves (first 500 s in the presence of 1 mM EGTA or 40 s in the presence of 100 μM CaCl_2_).

### In vitro motility assay

In vitro motility experiments were performed at 24–28 °C on glass surfaces silanized with trimethylchlorosilane (TMCS) as previously described [53, 54]. During a given experiment, the temperature was kept constant to within 1-2 ^°^C. Briefly, silanization was performed as follows; first glass cover-slips (60 × 24 mm^2^, #0, Menzel Gläser, Braunschweig, Germany) were cleaned with piranha solution (H_2_SO_4_ and 30% H_2_O_2_ at 7:3 ratio; *note that piranha solution is highly corrosive and acidic, which reacts violently with organic materials. Therefore, follow appropriate safety precautions*) at 80 °C for 5 min followed by sequential washing with H_2_O (thrice), methanol, acetone and chloroform. Cleaned glass coverslips were then functionalised with 5% TMCS in chloroform for 2 min and washed with chloroform. Thus functionalised surfaces were dried under dry N_2_ gas stream and stored under ambient conditions (Petri dishes sealed with parafilm) [54].

Motility assays were performed using the following buffer solutions: (1) Low ionic strength solution (LISS): 1 mM MgCl_2_, 10 mM MOPS, 0.1 mM K_2_EGTA, pH 7.4. (2) L65: LISS containing 50 mM KCl and 10 mM dithiothreitol (DTT). (3) A60 Assay solution containing 1 mM MgATP, 10 mM DTT, and 45 mM KCl added to LISS solution supplemented with an anti-bleaching mixture containing 3 mg/ml glucose, 0.1 mg/ml glucose oxidase, 0.02 mg/ml catalase, and an ATP regenerating system containing 2.5 mM creatine phosphate and 0.2 mg/ml creatine phosphokinase. Flow cells were assembled using double-sided adhesive tape to form a fluid chamber between a non-functionalized small cover slip (20 × 20 mm^2^) and a TMCS-functionalized large cover slip (60 × 24 mm^2^). In a typical motility assay, the flow cell was infused sequentially with the following solutions: (1) HMM (120 μg/ml) diluted in L65 for 5 min, (2) bovine serum albumin (BSA, 1 mg/ml) in L65 for 2 min, (3) Wash (L65, once) (4) F-actin (0.25 μg/ml) labelled with rhodamine-phalloidin in L65, (5) Wash (L65, thrice) (6) A60 (either cold, which is incubated for 2 min or pre-warmed). For gelsolin mediated F-actin severing experiments, gelsolin (1-10 nM) and calcium (details below) were added into the A60 assay solution, without changing the concentration of any other reagents from the values above. The fluorescence images of F-actin sliding on HMM were captured using an inverted fluorescence microscope (AxioObserver D1, Zeiss, Jena, Germany) with 63× planapochromat objective (Zeiss: 1.4 N.A). The image sequences were recorded using a digital CCD camera (C4742-95, Orca-ER, Hamamatsu Photonics, Hamamatsu, Japan) (Hamamatsu Photonics) or an EM-CCD camera (C9100, Hamamatsu Photonics) using HCImage software. The resolution of the recorded images, at an overall image size of 512 × 512 pixels, was 0.198 μm^2^/pixel. The frame rates used were 5 s^−1^. The image sequences were analysed using MatLab software (MatLab R2017a; MathWorks, Natick, MA) to obtain filament-sliding velocities [55, 56]. Filament lengths of sliding and stationary actin filaments were obtained from calibrated intensity data as described previously [57].

Assay solutions with different concentrations of calculated free Ca^2+^ were prepared for the experiments with and without gelsolin by modifying the above mentioned A60 solution as follows: 0 nM free Ca^2+^ (addition of 100 µM EGTA, no added CaCl_2_), 6.8 nM free Ca^2+^ (100 µM EGTA, 10 µM CaCl_2_), 1.1 µM free Ca^2+^ (100 µM EGTA, 97 µM CaCl_2_), 1.9 µM free Ca^2+^ (100 µM EGTA, 100 µM CaCl_2_). The free Ca^2+^ concentrations given below refer to the mixtures of EGTA and CaCl_2_ indicated in the previous sentence. Free calcium concentration was calculated by a Maxchelator program, version WEBMAXC STANDARD that is typically used to determine the free metal concentration present in the solution in the presence of chelators [58-60].

### TIRF assay

TIRF assay solution, pH 7.4 was prepared to contain (final concentrations): 2 mM Trolox, 2 mM Cyclooctatetraene (COT), 2 mM 4-Nitrobenzyl alcohol (NBA), 10 mM DTT, 45 mM KCl, 7.2 mg/ml glucose, 3 U/ml POX, 0.01 mg/ml catalase, 2.5 mM creatine phosphate and 0.2 mg/ml creatine phosphokinase in LISS [61]. Initially, 100 mM Trolox stock was prepared in methanol, followed by dilution in LISS, subsequent filtering through a 0.2 µm filter and exposure to UV light (254 nm, 120,000 µJ/cm^2^) for 15 min to form Trolox-Quinone. The Trolox-Trolox/Trolox-Quinone mixture prepared in LISS (TT/TQ-LISS) was degassed before use. For further details, please see [61].

The TIRF assay was performed by first adsorbing HMM (12 µg/ml) onto TMCS-derivatized glass surfaces (5 min). Subsequently, the flow-cell was infused sequentially with 1 mg/ml non-fluorescent BSA (2 min), wash buffer L65, rhodamine-phalloidin labelled F-actin (5 nM, 2 min) and gelsolin (5 nM, labelled with Alexa-647) with varying concentrations of free calcium (see above) as desired. All TIRF assay experiments were performed at a stable temperature (23 ± 1 °C), using an objective heater [61]. Time-lapse movies were acquired with an exposure time of 50 ms. An objective-based TIRF illumination system was built in house using a Nikon TIRF 60X objective (NA=1.49), a Nikon Eclipse TE300 inverted microscope, an Andor iXon Ultra 897 EMCCD camera and a 642 nm diode laser for illumination as described elsewhere [61].

The simultaneous addition of NBA, COT and Trolox, to improve dye photo-physical properties in the TIRF assay, reduced the actin filament sliding velocity whereas Trolox alone had negligible effects (compare Fig. S1A to Fig. S1B). Importantly, however, the effects of gelsolin on velocity were similar whether only Trolox or all compounds were present (compare Fig. S1A to Fig. S1B), justifying the use of all components to ensure optimal image quality.

## Statistical analysis

Data were analysed using MatLab software as described above and the subsequent non-linear and linear curve fittings and statistical analyses were performed using Graphpad Prism (version 8.2.1, Graphpad software, CA). Unless otherwise stated, data are given as mean ± 95% confidence interval (CI).

## Results and Discussion

### Gelsolin accelerates actin depolymerisation in bulk assays

The actin severing activity of gelsolin is complex. A Ca^2+^ concentration in the micromolar range (∼0.1-5 µM) is typically required to modify gelsolin to its active conformation (closed to open) that is competent for F-actin severing and barbed end capping [51, 62-64]. However, other experimental conditions (e.g. concentration of reagents used in the assay in addition to Ca^2+^, such as gelsolin itself, actin, myosin and ATP) may influence the gelsolin binding to actin filament and the gelsolin-mediated filament splitting rates.

To investigate the basal F-actin severing activity of the gelsolin preparations used in the present study under standardized conditions, we first checked the disassembly of actin filaments in-solution in the presence of EGTA (1 mM) or CaCl_2_ (100 μM) (Fig. 1). These conditions correspond to free Ca^2+^ concentrations (calculated) of zero (0) and 95.3 µM respectively. We used F-actin formed upon polymerization of 50% pyrene labelled G-actin followed by dilution to a final concentration of 20 nM, below the barbed end critical concentration. These conditions favour the monitoring of spontaneous monomer dissociation. Filament disassembly kinetics was monitored by recording the pyrenyl emission as a function of time. In a first set of experiments, we found that the rate of spontaneous actin depolymerisation is very low in the presence of EGTA, whereas addition of CaCl_2_ slightly favours the monomeric form of actin even in the absence of gelsolin (Fig. 1A). This may be attributed to the effects of the divalent cation on the mechanical properties of F-actin [65, 66].

**Fig.1.**
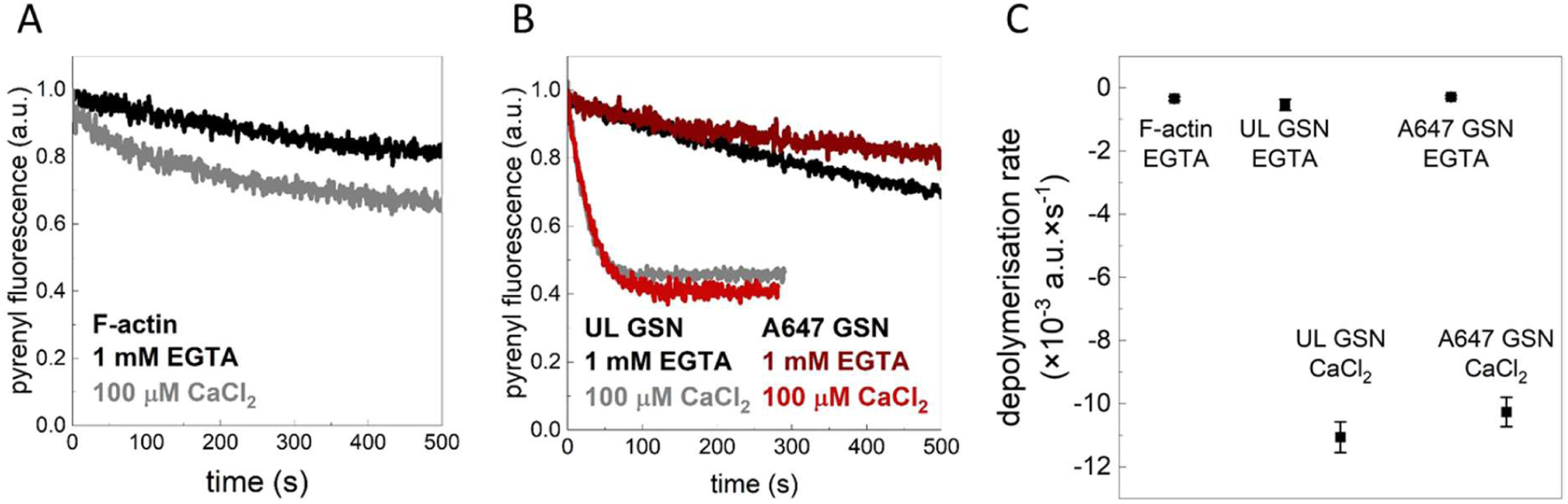
Ca2+ enhances the actin filament severing activity of gelsolin. For the depolymerisation assays, actin was polymerized and diluted in polymerisation buffer to a final concentration of 20 nM to induce spontaneous depolymerisation. Gelsolin (GSN) was used at 5 nM final concentration along with 100 µM Ca^2+^ (free Ca^2+^ = 95.3 µM) or 1 mM EGTA (0 µM free Ca^2+^). A) Spontaneous actin disassembly in the presence of either Ca^2+^ (100 µM) or EGTA (1 mM) but in the absence of GSN. Data are presented as the average of two independent measurements at each condition. B) Actin disassembly in the presence of gelsolin (5 nM) (unlabelled; UL or Alexa-647 labelled; A647) and in the presence of either Ca^2+^ (100 µM) or EGTA (1 mM). Data are presented as the average of 2 (A647 GSN) and 4 (UL GSN) independent measurements. C) Depolymerisation rate (negative) derived from linear fitting of the pyrenyl transients. Mean ± SD, n = 2 – 4. Note negligible effects of gelsolin on depolymerisation rate in the absence of Ca^2+^ but a substantial reduction at 100 µM added Ca^2+^.

Next, we investigated the F-actin severing activity of gelsolin using two different gelsolin batches, where gelsolin was either unlabelled or Alexa-647 labelled (Fig. 1B). The results from these assays, summarized in Fig. 1C, demonstrate that in the absence of Ca^2+^ (1 mM EGTA) actin disassembly was not significantly influenced by addition of gelsolin (5 nM) at shorter timescales (< 200 s) (Fig. 1B-C). However, a slightly faster rate than that for spontaneous disassembly was detected in the presence of unlabelled gelsolin (GSN_EGTA_/F-actin_EGTA_ ∼ 1.58-fold) as compared to the labelled protein, an effect that became apparent at longer times. As expected, the disassembly rate was appreciably increased (GSN_CaCl2_/GSN_EGTA_ ∼ 20-35-fold) upon changing to excess CaCl_2_ (free [Ca^2+^] ∼95.3 µM) whether unlabelled or Alexa-647 labelled gelsolin was used (Fig. 1B and Fig. 1C). Thus, the disassembly rates both in the absence and in the excess of Ca^2+^ was similar for Alexa-647 labelled and unlabelled gelsolin and both proteins showed strong calcium dependent severing activity.

### Cooperativity between gelsolin mediated and myosin mediated effects on actin filaments

To the best of our knowledge, the severing activity of gelsolin during an in vitro motility assay has not previously been reported. Here we performed studies using this assay to elucidate the interactions between gelsolin mediated F-actin severing and actin-myosin motor activity.

We tested different concentrations of gelsolin in the nanomolar range (0.75-1 nM) [67] with a concentration of F-actin corresponding to subunit concentrations in the range of 3-5 nM. The studies were performed in assay solution A60 containing 100 µM EGTA under low and high [Ca^2+^], defined as ∼6.8 nM and 1.9 µM free Ca^2+^, respectively. First, we noted that the number of filaments in the in vitro motility assay increased with time even in the absence of gelsolin due to fragmentation by myosin-induced forces (Fig. 2, A60_control). If the gelsolin concentration was in the range of 2-10 nM at free [Ca^2+^] of 1.9 µM, the actin filaments (added to flow cell at 3 nM; monomer concentration) disappeared rapidly (Fig. S2D), leaving virtually no observable filaments 30 s after addition of the ATP-containing assay solution (Fig. S2). We attribute this effect to rapid severing and disassembly of F-actin [68, 69]. We also tested gelsolin concentrations ‡1 nM (0.75-1.0 nM) at high calcium concentration of 1.9 µM free Ca^2+^.

**Fig.2.**
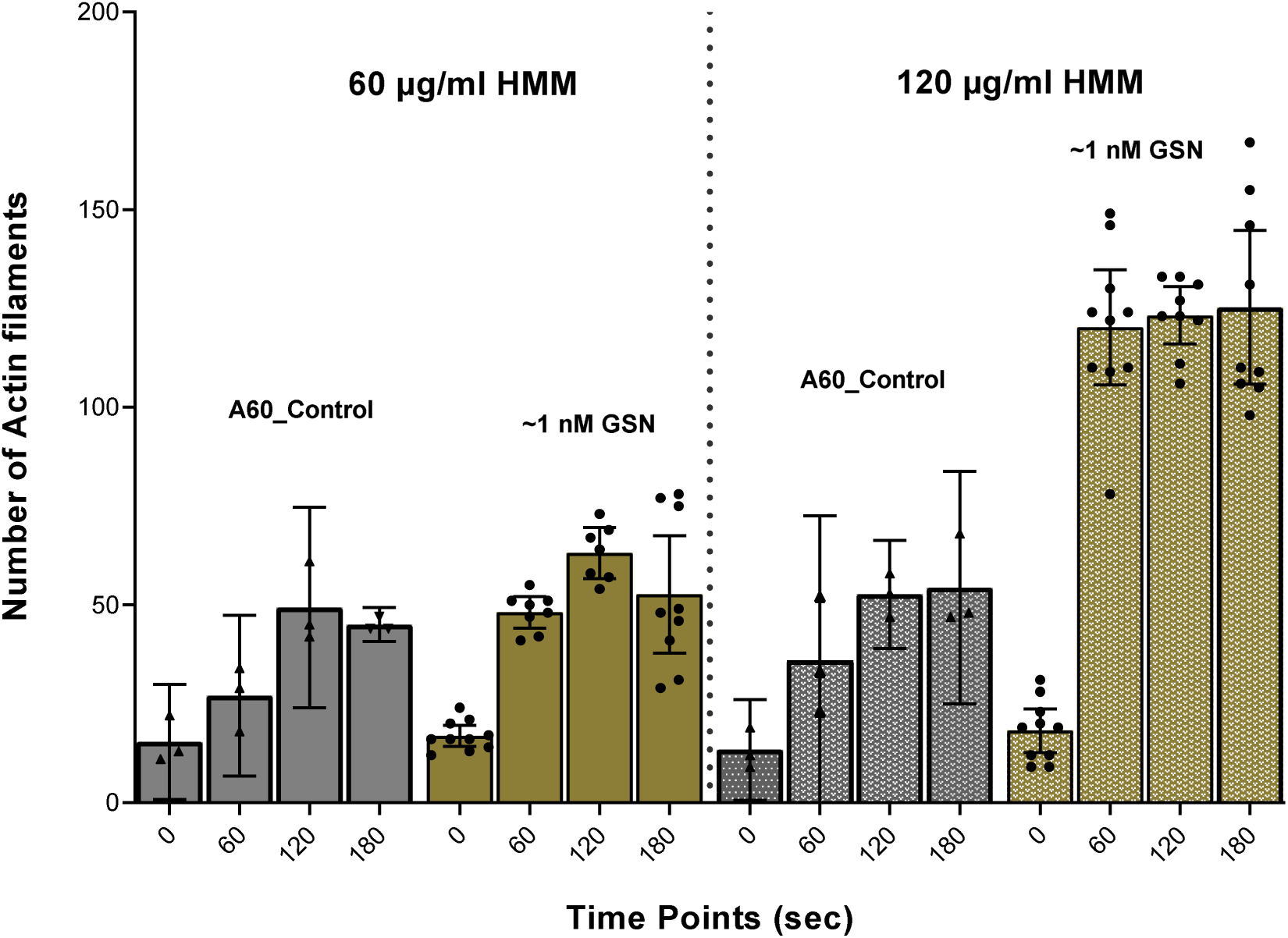
Number of actin filaments in the in vitro motility assay at different time points after addition of MgATP containing assay solution including 1.9 µM free Ca^2+^ with or without gelsolin (GSN; ∼1 nM), and different surface incubation concentrations of HMM (60 or 120 µg/ml). Data shown as mean ± 95% CI. Temperature: 25-27 °C. Data from individual experiments superimposed on bars representing mean values.

Under these conditions, using HMM incubation concentration of 60 µg/ml, we observed increased (up to 2-fold) severing of actin filaments in the in vitro motility assay in the presence of gelsolin as compared to its absence, as quantified by an increase in the number of actin filaments at 60 s (Fig. 2). The HMM incubation concentration of 60 µg/ml that was used, is expected to give less than saturating density of HMM on the motility assay surface (e.g. [57] [70]; supplemental information of latter paper). Remarkably, with a higher HMM incubation concentration (120 µg/ml), expected to saturate the surface [57], there was a 5-7-fold increase in the number of actin filaments within 60 s in the presence of gelsolin as compared to control samples (Fig. 2). The increase in the filament number with time was significantly higher than in the absence of gelsolin as indicated by non-overlapping 95% CIs. Moreover, noticeably, the increased HMM surface density had negligible effects on the motor induced severing in the absence of gelsolin. The marked difference between the effects of increased HMM density on motor induced severing in the presence and absence of gelsolin, suggests cooperative effects between gelsolin and motor function in this regard.

We also studied the effects of varying HMM density on gelsolin-mediated fragmentation at low (6.8 nM) free [Ca^2+^] as illustrated in Fig. 3, where reduced fragmentation rate is observed for reduced HMM incubation concentration in the range 120 – 60 µg/ml. It can be seen that the increase in the number of filaments with time is somewhat less at 5 nM gelsolin and low [Ca^2+^] than at 1 nM gelsolin and high [Ca^2+^] (compare Fig. 2 and Fig. 3). However, importantly, in both cases, the number of filaments increased to greater extent in the presence of gelsolin during the first 60 s and the effect of gelsolin was greater at higher HMM surface density. The findings suggest that gelsolin binds to actin filaments at both low and high [Ca^2+^]. Furthermore, also gelsolin bound to actin at low [Ca^2+^] potentiates fragmentation of the filaments due to forces produced by myosin motor activity. To summarize, the results in Fig. 3 suggest that, also at low [Ca^2+^], gelsolin addition noticeably increases the actin filament severing with increased HMM density whereas gelsolin has negligible effects on the motor induced severing at the lowest HMM density. These findings are consistent with cooperative effects between motor induced and gelsolin induced severing also at low [Ca^2+^].

**Fig.3.**
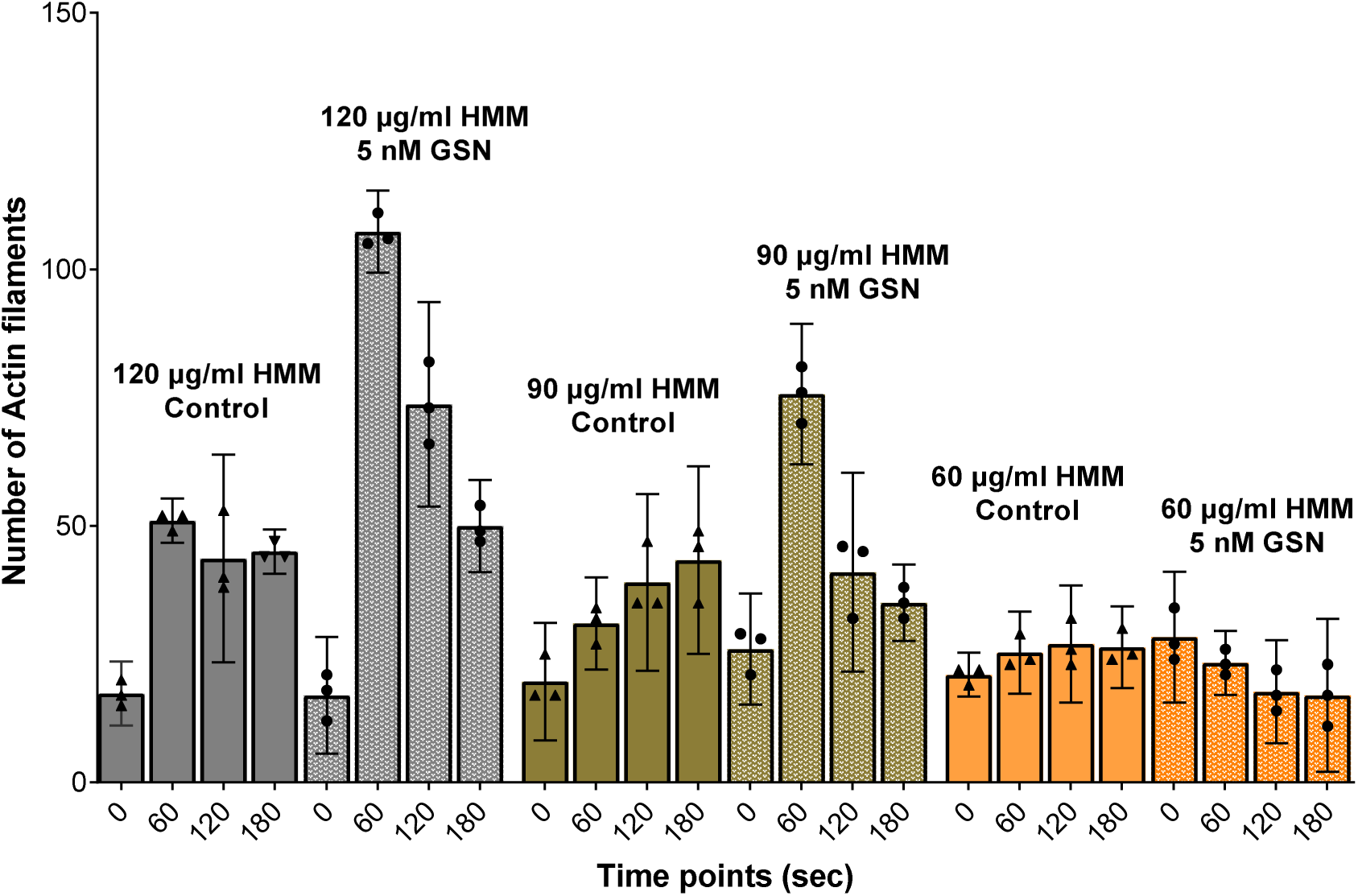
Number of actin filaments in the in vitro motility assay at different time points after incubation with HMM at different concentrations; 60-120 µg/ml, ∼6.8 nM free Ca^2+^. Note, we attribute the decrease in the number of filaments at the later time points to detachment of filaments from surface and difficulty to observe short filaments due to weak total fluorescence. Data shown as mean ± 95% CI. Temperature: 25-27 °C. Data from individual experiments superimposed on bars representing mean values.

To further investigate the interactions between gelsolin, actin and myosin (HMM) in the presence of MgATP at low [Ca^2+^], we analysed gelsolin-induced changes in the filament sliding velocity in the in vitro motility assay at a free [Ca^2+^] of ∼6.8 nM. Generally, the average velocity decreased with increased gelsolin concentration in the range of 1-10 nM. To get insight into the mechanistic basis for this effect, we performed detailed analyses of the dependence of the sliding velocity on filament length using in vitro motility assays at HMM incubation concentration of 120 µg/ml. In these analyses, we compared 5-10 nM gelsolin to conditions without gelsolin. The results (Fig. 4) show that the presence of gelsolin reduces average velocity by: a) reduced velocity of the longest filaments studied (rightmost open squares in Fig. 4A), b) greater reduction in sliding velocity at filament lengths below about 2 µm (leftmost open squares in Fig. 4A) and c) reduction in the average length of the studied filaments.

**Fig.4.**
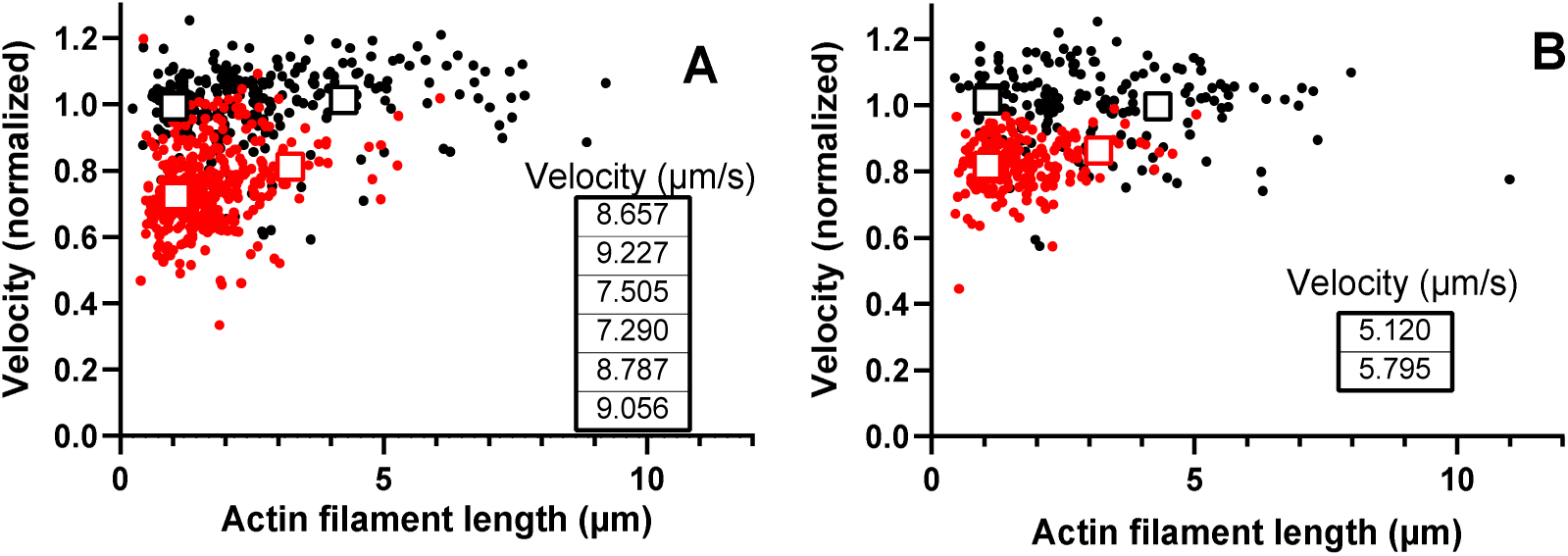
Effects of gelsolin (5-10 nM) in the presence of free Ca2+ (∼6.8 nM) on HMM-induced F-actin sliding velocity vs filament length in the in vitro motility assay. A) HMM incubation concentration 120 µg/ml in the absence (black dots) and presence (red dots) of gelsolin. Average sliding velocities ± 95% CI (within size of the symbol) shown as open squares for control (black) and gelsolin (red) conditions for filament lengths < 1.5 µm (left squares) and > 2.5 µm (right squares), respectively. Data points located at the average filament lengths in the respective range. Data from 6 experiments. Before plotting, data in each individual experiment (30 – 80 filaments) were normalized to the average absolute velocity in the absence of gelsolin (see inserted table). B) HMM-induced F-actin sliding velocity with HMM added at 60 µg/ml in the absence (black dots) and presence (red dots) of gelsolin. Average sliding velocities ± 95% CI (within size of the symbol) shown as open squares control (black) and gelsolin (red). Data from 2 different experiments, plotted as in A, with the average velocity in each individual experiment (61 and 106 filaments) in the absence of gelsolin given in inserted table. Temperature: 25-27 °C.

We attribute a somewhat greater reduction in velocity for short compared to long filaments to either of two mechanisms: i) gelsolin induced structural changes propagate only short distance along the actin filament or ii) gelsolin changes the actin filament structure to reduce the myosin association rate constant along the entire length of all filaments thereby reducing the duty ratio. In the latter case one would expect greater reducing effect on velocity for short filaments because such filaments interact with fewer HMM molecules. If this is the key mechanism, one would also expect greater reduction in velocity (particularly for short filaments) at low (e.g. 60 µg/ml), compared to high (e.g. 120 µg/ml) HMM surface density. This was not observed (Fig. 4B), arguing for the alternative idea (i) of more extensive gelsolin induced structural changes in F-actin close to the gelsolin binding site at the barbed end compared to more distant locations along the filament. This is more likely than the alternative mechanism to cause similar effects on the velocity-length plot at low and high HMM density (Fig. 4A-B).

### On the possibility of gelsolin mediated displacement of phalloidin from the actin filaments

In vitro motility assay experiments performed in the presence of gelsolin resulted in the eventual loss of observable rhodamine phalloidin labelled actin filaments from the motility assay surface as noted by tendency for a decrease in the number of filaments at times > 60 s in Fig. 3 and Fig. S2C, D and S3B. These effects could be attributed to either of the following effects: 1. production of very short filaments either detaching from the surface or exhibiting too faint fluorescence (further aggravated by photobleaching) to be observed in epi-fluorescence microscopy or 2. gelsolin induced displacement of phalloidin from actin filaments [71].

To investigate the latter possibility, we used covalently linked NHS-rhodamine instead of phalloidin-rhodamine for fluorescence labelling of the actin filaments. NHS-rhodamine labelled actin filaments showed relatively weak fluorescence signal compared to phalloidin-rhodamine. In control motility assays, both actin filaments labelled by NHS-rhodamine and phalloidin-rhodamine behaved in apparently similar manner with long-term and durable motility (Fig. S3A, S3C). Motility assays performed with NHS-rhodamine labelled filaments in the presence of gelsolin (5 nM) and low calcium, (free Ca^2+^ ∼6.8 nM) resulted in slightly faster disappearance of the actin filaments (Fig. S3D) than in the case with phalloidin-rhodamine (Fig. S3B). Therefore, these results suggest that gelsolin-based displacement of phalloidin-rhodamine was not the basis for a reduced number of filaments with time in the presence of gelsolin. Rather, we favour the alternative explanation based on production of very small filament fragments that are either not observable due to faint fluorescence or that detach from the surface. This suggests an underestimation of the increase in the number of filaments with time in the in vitro motility assay in the presence of gelsolin (Figs. 2, 3) and thus underestimation of the severing efficiency cooperatively attributed to gelsolin and myosin induced forces.

### One-to-one relation and co-localisation between gelsolin binding and severing in the absence of motor forces

We show above that high gelsolin and calcium concentrations induce rapid severing of the actin filaments under in vitro motility assay conditions. For practical purposes, to be able to follow the time course, we had to limit either the gelsolin concentration or the Ca^2+^ concentration.

In order to study severing at high concentrations of both gelsolin (5 nM) and free Ca^2+^ (∼1.9 µM), we turned to total internal reflection fluorescence (TIRF) microscopy to enable observation down to single fluorophores by limiting illumination to a ∼100 nm thick layer above the surface. Furthermore, to maintain high resolution and prevent detachment of very short actin filaments from the surface, as may occur in an in vitro motility assay (see above), the filaments were first kept stationary on the surface by binding to HMM (added at 12 µg/ml) without any added MgATP in a TIRF assay solution (see Materials and Methods). This facilitates observation of both individual actin filaments and Alexa-647 labelled gelsolin and therefore of the gelsolin-binding and the associated severing pattern along F-actin. Using these conditions, we first noted a slow gelsolin mediated fragmentation (half-life ∼5 min) of the filaments in the absence of Ca^2+^ (Fig. 5A), in agreement with bulk solution fluorescence spectroscopy data (Fig. 1). By increasing the gelsolin concentration to 5 nM and the free [Ca^2+^] to ∼1.9 µM the severing became very fast and, within 15 seconds, there were no visible actin filaments left on the surface (Fig. 5B and 5C) with similar behaviour using Alexa-647 labelled and non-fluorescent gelsolin.

**Fig.5.**
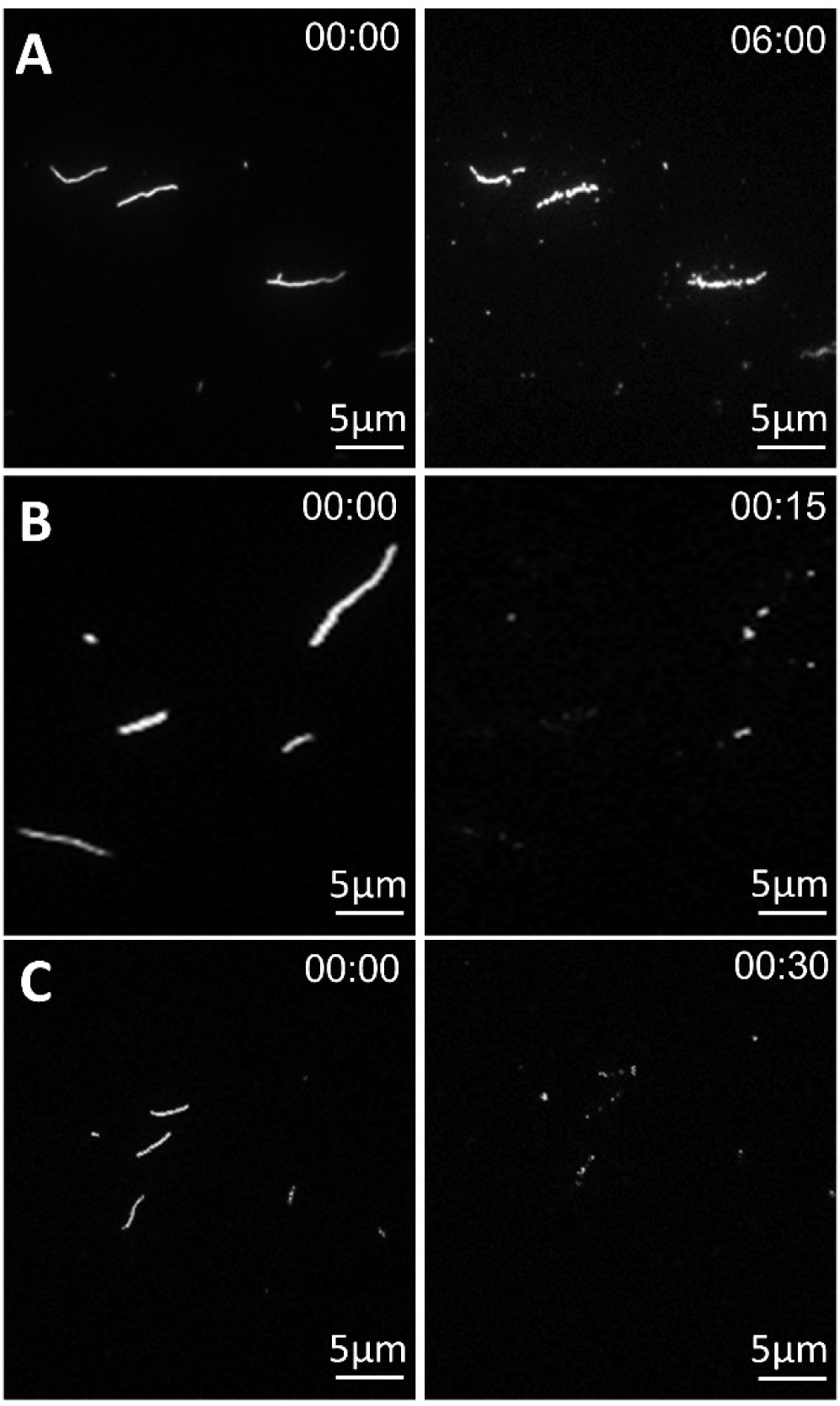
TIRF microscopy images showing F-actin severing activity in the presence of 5 nM gelsolin. A) Unlabelled gelsolin without calcium, B) Unlabelled gelsolin with free Ca^2+^ ∼1.9 µM) and C) Alexa-647 labelled gelsolin and free Ca^2+^ ∼1.9 µM. HMM = 12 μg/ml. Assay solution was added without MgATP, resulting in a rigor actomyosin state. Time points are represented as minutes:seconds.

There is evidence (e.g. from Figs. 2-4) and data in the literature (e.g. [46]) that gelsolin, when bound to an actin filament, produces long-distance changes in filament structure. We therefore asked if gelsolin severing in the absence of HMM driven motion only occurs at the gelsolin binding site or if severing could occur at distant sites along the filament. To test this idea, we used Alexa-647 labelled gelsolin and TIRF microscopy based single molecule studies. As shown above we observed similar severing activity of the unmodified and the Alexa-647 labelled gelsolin whether using bulk depolymerisation assays (Fig. 1) or TIRF microscopy-based observation of the filaments on a surface (Fig. 5).

In the absence of Ca^2+^, we observed only limited binding of Alexa-647 labelled gelsolin molecules along the length of the actin filament and very limited filament severing activity (Fig. 6A; however, cf. Fig. 5). Of the 10 filaments observed, there were on average 1.7 gelsolin attachments with 0.5 cuts per filament during 9 min observation time. Addition of free Ca^2+^ at 0.47 µM (Fig. 6B) was ineffective in inducing actin filament severing in the absence of motor induced forces. However, at free Ca^2+^ of ∼1.9 µM, rapid severing of actin filaments was observed, similar to the results obtained using unlabelled gelsolin (Fig. 5). Our TIRF microscopy to observe Alexa-647 fluorescence (Fig. 6C) showed binding of multiple gelsolins along the length of the actin filaments and filament severing was primarily observed at the points of observed gelsolin attachment. With 49 cuts observed on 9 filaments Alexa-647 labelled gelsolin was bound at 34 of the cuts. This gives a #gelsolin/#cuts ratio of 0.69 comparable to the Alexa-647/gelsolin labelling ratio of 0.67. These results support the view that gelsolin-mediated filament severing of stationary filaments is a local event due to conformational changes at the point of gelsolin binding. The results do not support ideas that severing takes place also at sites distant from the gelsolin binding site if the filaments are not propelled by HMM.

**Fig.6.**
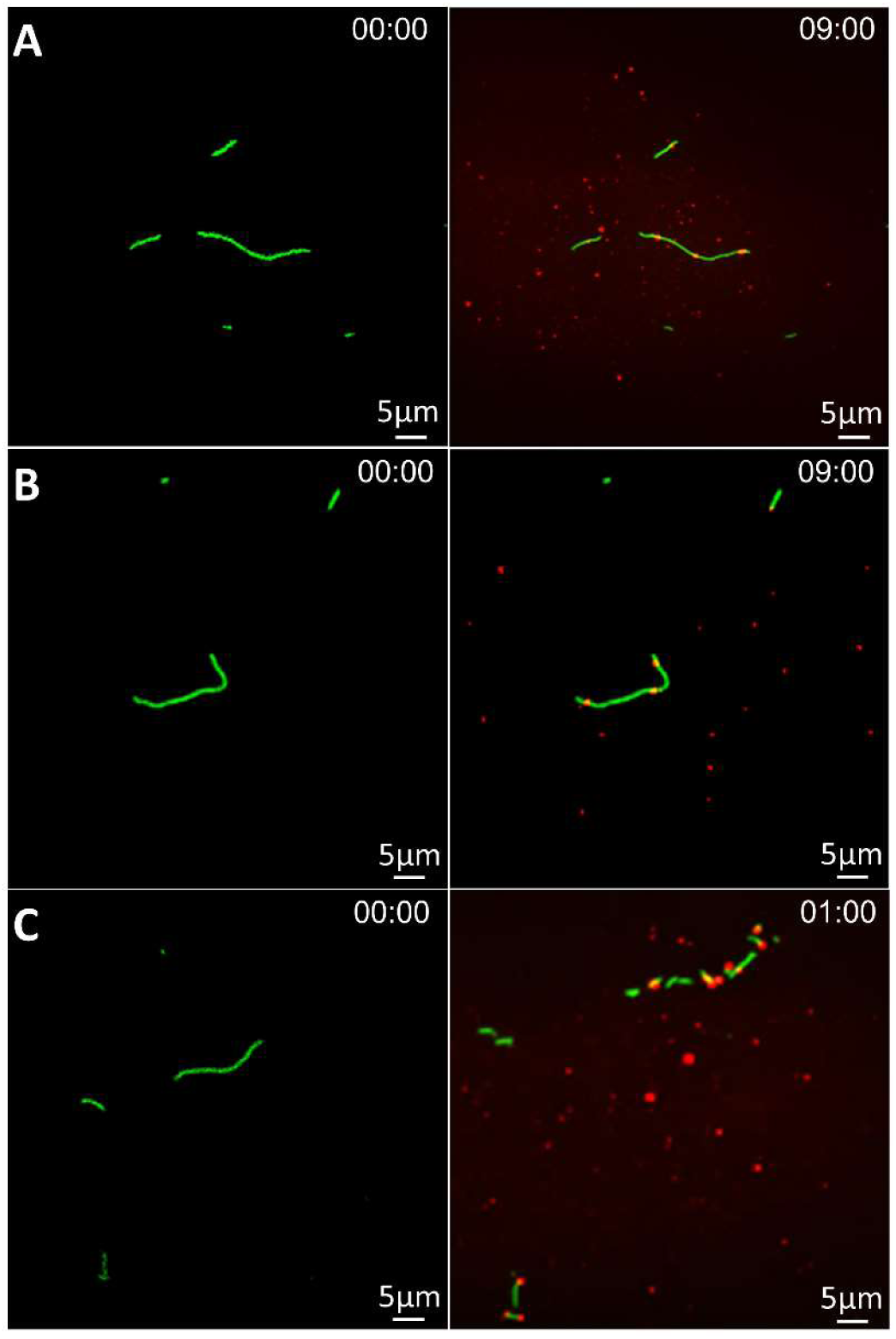
TIRF microscopy images showing gelsolin binding and or severing. F-actin severing activity in the presence of 5 nM Alexa-647 gelsolin. A) No calcium B) free Ca^2+^∼0.47 µM; C) free Ca^2+^ ∼1.9 µM. HMM = 12 µg/ml. Assay solution was added without MgATP resulting in a rigor actomyosin state. Time points are represented in minutes:seconds.

### Gelsolin-binding to actin and myosin during in vitro motility assays

The magnitude of the increase in number of actin filaments and the rate of this increase were appreciably higher during in vitro motility assays at 5 nM gelsolin and low (∼6.8 nM) free [Ca^2+^] (compared to no gelsolin; Figs. 3-4) than for stationary filaments (Figs. 5-6). The structural changes induced by the gelsolin binding at low [Ca^2+^] thus seem sufficient to harness the motor induced forces more effectively to produce filament severing. The gelsolin mediated severing of stationary filaments required higher free [Ca^2+^] and was characterized by a sharp threshold in [Ca^2+^] from 1.1 µM to 1.9 µM free Ca^2+^ (5 nM gelsolin). Above this threshold, further increase in Ca^2+^ concentration caused apparently complete and rapid fragmentation with disappearance of the filaments (consistent with excess gelsolin concentration above the actin concentration on monomer level). Furthermore, gelsolin mediated actin severing also revealed capping of one end of pre-existing actin filament by labelled gelsolin (Fig. 6).

To gain more insight into the gelsolin effects in the in vitro motility assay at low [Ca^2+^] (Figs. 3-4) we performed in vitro motility assays under TIRF illumination in the presence of 5 nM Alexa-647 labelled gelsolin. The results (Fig. 7) confirmed gelsolin binding to a substantial fraction of the HMM propelled actin filaments (36.5%, n = 282 filaments from 10 images). Furthermore, interestingly, only those actin filaments that had bound gelsolin showed reduced velocity (Fig. S1). This corroborates the idea that the reduction in velocity is attributed to the bound gelsolin and it is consistent with gelsolin-induced structural changes along the length of the filament. Considering the labelling efficiency of gelsolin with Alexa-647 (∼ 67%) the observed 37% gelsolin labelling in Fig. 7 suggests that approximately 55% of all sliding filaments have bound gelsolin. Unfortunately, the locations of these gelsolin molecules along the filaments were not possible to determine using our set-up due to limited spatial resolution. However, it is interesting to note that the results with a little more than two-fold doubling of the actin filament severing upon addition of gelsolin at the highest motor density in Fig. 3, together with the gelsolin binding by 55% of the filaments (Fig. 7) is consistent with gelsolin capping at the barbed end, i.e. local cutting of the filament close to the gelsolin molecule. Similarly, the data in Fig. 3 at lower motor densities would then suggest that the motor induced forces under these conditions are insufficient to achieve cutting, as was the case with thermal fluctuations (Fig. 6).

**Fig.7.**
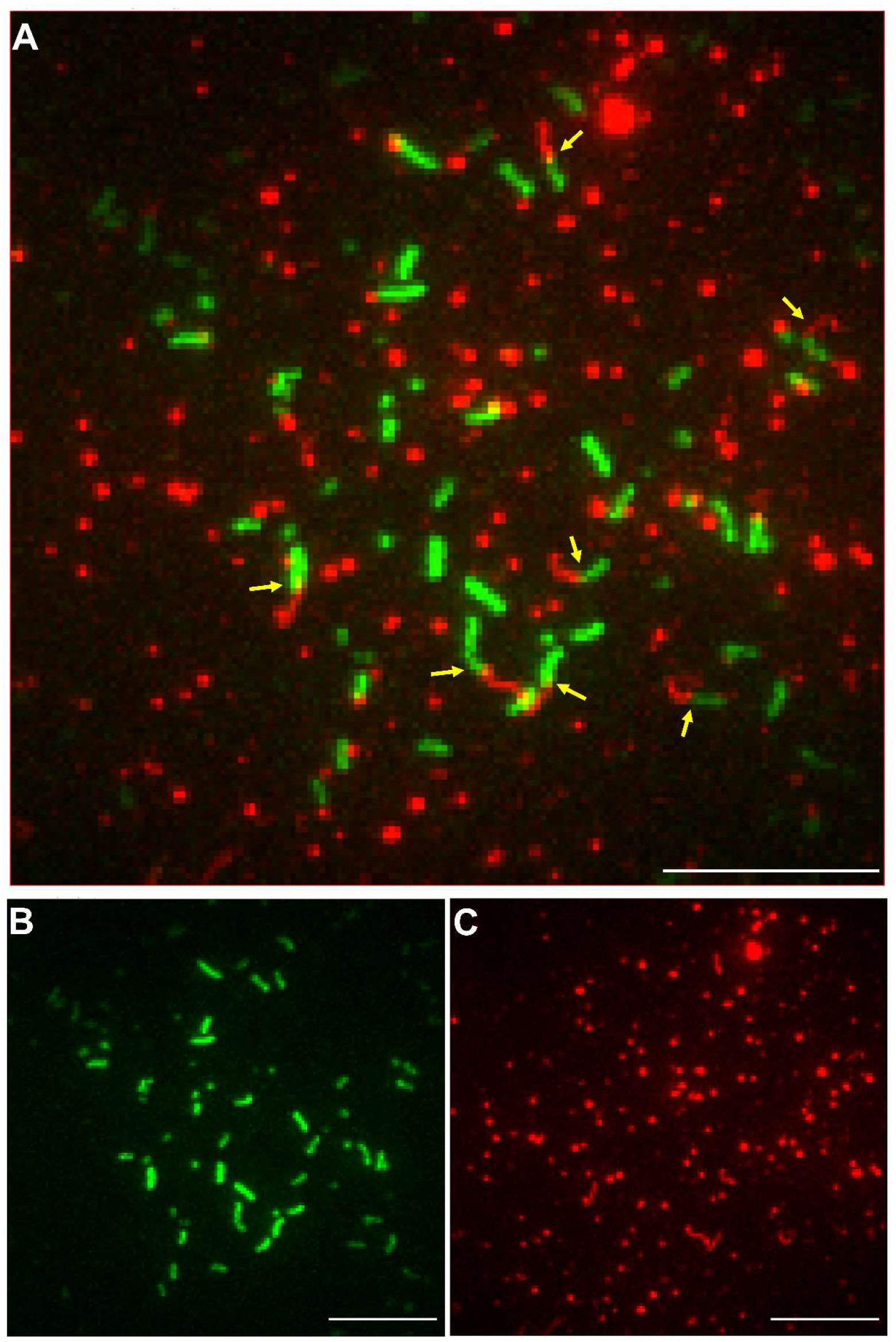
Maximum Z-projection images of pseudocoloured actin filaments (green, Alexa488 phalloidin labelled) and gelsolin (red, Alexa647 labelled). A) Z-projection of both actin filaments and gelsolin (30+30 frames at 20 s^-1^ frame rate). For the first 30 frames, filter cube to observe Alexa488 phalloidin was used, followed by switch to filter set to observe Alexa647 labelled gelsolin. Yellow arrows denote motile filaments with gelsolin attached as verified by visible filament trajectories both before and after change of filter cube. B) Z-projection of actin filaments (30 frames); C) Z-projection of gelsolin bound to filaments and non-specifically to the surface. Images were acquired using individual channel filters by TIRF microscopy and colocalised using ImageJ (Ver. 1.53a) (image scale = 10 µm)

It is also of interest to investigate the degree of gelsolin-binding to the in vitro motility assay surface in the absence of actin because non-specific surface binding of gelsolin may produce frictional forces between gelsolin labelled filaments and the underlying surface, contributing to a reduced sliding velocity (cf. Fig. 4). We performed such studies by observing single molecule fluorescence of Alexa-647 labelled gelsolin (Fig. S4) and found that the non-specific surface adsorption of gelsolin was markedly increased at high compared to low [Ca^2+^]. This is consistent with a more flexible gelsolin structure at micromolar Ca^2+^, because “soft proteins” are known [72] to show an increased propensity for surface adsorption. Furthermore, we noted that the surface adsorption was similar in the presence and absence of HMM incubation arguing for non-specific binding to the underlying surface rather than to HMM. The limited non-specific binding of gelsolin at Ca^2+^ concentrations (Fig. S4B) similar to those used in the experiments in Fig. 4 argues against major contributions from gelsolin-surface friction to the reduction in sliding velocity of gelsolin-bound filaments. That gelsolin, immobilized to the underlying surface, should interact with actin is unlikely because HMM at saturating density holds actin ∼50 nm away from the surface, much greater than the average gelsolin diameter (∼10 nm) [63, 73]. Furthermore, importantly, the filament length dependence and the HMM surface density dependence of the gelsolin induced reduction in sliding velocity (Fig. 4) argue against a reduction caused by frictional forces due to interactions between HMM (or the surface) and the actin-bound gelsolin. Any such effects would have been negligible for long filaments due to increasing motor induced forces that overcome the friction and the effects would have been more substantial at low HMM density because of lower overcoming motor forces. Neither of these effects were observed in Fig. 4.

### Overall mechanistic interpretations of the data

The results reflect a complex interplay between actin, HMM, [ATP], [Ca^2+^] and gelsolin. Myosin generates contractile, extensile, bending and torsional forces [74-76] with shearing, buckling and eventually severing of the actin filaments [26] even in the absence of actin-binding proteins such as cofilin and gelsolin. Myosin-driven actin filament fragmentation in the in vitro motility assay experiments occurs in the presence of nearly physiological millimolar concentrations of ATP [26]. The motility experiments suggest that the gelsolin-mediated F-actin severing cooperates with the severing due to myosin motor function, in the sense that a given gelsolin concentration caused more extensive severing (greater increase in the number of filaments) at increased HMM surface density. Conversely, increasing concentrations of gelsolin (even at low Ca^2+^-concentrations) led to increased actin filament fragmentation at a given HMM density (Fig. 3). These effects can be accommodated by local effects of gelsolin at its binding site to the actin filament and does not necessarily rely on gelsolin induced long-range allosteric changes in filament structure as suggested previously [74, 77]. However, the effects of gelsolin binding on the sliding velocity at different filament lengths are difficult to explain without such long-range changes along the actin filament, previously suggested based on spectroscopic [29, 47] and electron microscopic [46, 78] evidence. Types of long-range changes that have been observed include changes in orientation (by 10°) and a (threefold) decrease in torsional rigidity and hence effects on the helical structure and torsional flexibility throughout the whole actin filament [29, 77]. However, our results showing filament velocity as a function of filament length (Fig. 4) suggest that the structural changes are more prominent at short distances from the gelsolin binding site at the barbed end. Strikingly, our results suggest that structural changes along the actin filament upon gelsolin binding are produced both at micromolar and nanomolar free [Ca^2+^]. This is partially consistent with the results shown in [48], where it was proposed that the degree of calcium binding to gelsolin at very low (∼10 to 50 nM) [Ca^2+^] causes structural change that unlatches the closed structure of gelsolin [48]. The necessity for simultaneous myosin motor activity to effectively harness the gelsolin effects at low [Ca^2+^] follows from results in Figs 3, 1 and 6. Thus, appreciably increased filament severing was observed upon gelsolin addition during myosin induced sliding at high HMM density (Fig. 3). In contrast, very limited severing was observed in the absence of such forces (Fig. 1, Fig. 6) or with low forces of this type (low HMM densities in Fig. 3).

In the in vitro motility and TIRF microscopy assays, the actin filaments were generally labelled with rhodamine-phalloidin whereas in the bulk depolymerisation assays pyrene labelled actin was used. Some previous studies have emphasized the inhibitory role of phalloidin in depolymerisation of actin filaments [9]. However, phalloidin did not noticeably interfere with the gelsolin-mediated severing of actin filaments, in agreement with other previous reports [79, 80]. In effect, the use of high concentration of gelsolin and Ca^2+^ resulted in extremely rapid severing of the phalloidin labelled filaments both in motility assays and in TIRF microscopy based single molecule studies.

The results of TIRF based motility assays (Figs. 7, and S1) broadly corroborated the results of our regular motility assays (Figs. 2-4) using non-labelled gelsolin, both showing an appreciable reduction of the average sliding velocity. When gelsolin was used in the presence of high calcium, actin filaments underwent rapid severing and hence quickly disappeared. However, in the presence of low calcium, we could clearly observe the binding of gelsolin to the moving actin filaments in the motility assay, with a clear pattern of gelsolin moving along the path of the actin filaments (Fig. 7). Furthermore, actin filaments that were confirmed by the Alexa-647 co-localisation to have bound gelsolin experienced reduced average sliding velocities compared to non-labelled filaments, consistent with conformational changes along the actin filament when gelsolin is bound [81]. Alternative explanations for the effects of gelsolin on sliding velocity (Fig. 4), e.g. due to gelsolin-surface interactions are inconsistent with evidence presented above.

In conclusion, we have found strong evidence for cooperative effects between gelsolin induced changes in the actin filaments and changes due to myosin motor activity in the following respects. The gelsolin induced changes in filament structure 1. increase the susceptibility of the filament, locally at the gelsolin binding site, to myosin induced fragmenting forces and 2. alter the actin-myosin interactions along the actin filament to reduce the myosin induced sliding velocity for actin filaments of varying lengths but particularly for short filaments. Altogether the results argue for long-range structural changes in the actin filaments that, however, seem to be more prominent at shorter distances from the gelsolin capped barbed end. This is suggested by circumstantial evidence for local cutting at the gelsolin binding site and direct evidence for a reduction in sliding velocity of HMM propelled actin filaments with bound gelsolin. The results also suggest that extremely effective severing of actin filaments may be achieved by combined actions of gelsolin and myosin of possible physiological relevance under certain circumstances [82, 83].

## Acknowledgements

This work was funded by EU Horizon 2020 Research and Innovation Framework Programme of the European Union (grant agreement 732482) (Future and Emerging Technologies; Bio4comp), The Swedish Research Council (grant numbers 2015-05290 and 2019-03456), and The Faculty of Health and Life Sciences at The Linnaeus University, Sweden. We also acknowledge support by the GINOP-2.3.2-15-2016-00049 and EFOP-3.6.2-16-2017-00006 grants, as well as by PTE ÁOK-KA-2019-50 (to TH). We are grateful to Robert C. Robinson for providing the plasmid of human gelsolin.

